## Supplementary Materials for "Fat and carbohydrate interact to potentiate food reward in healthy weight but not in overweight or obesity"

**Table S1.** Descriptive statistics for subjective ratings and willingness to pay (WTP) for foods across the three macronutrient categories from participants with healthy weight (HW; n = 30).

| **Subjective Variable**  (Units or scale range) | **Carbohydrate**  M ± SD, *Range* | **Fat**  M ± SD, *Range* | **Combo**  M ± SD, *Range* |
| --- | --- | --- | --- |
| Liking (-100 to +100%) | 18.6 ± 31.4, *-100 to +100* | 17.3 ± 32.0, *-100 to +83.0* | 20.3 ± 36.1, *-100 to +89.3* |
| Familiarity (0-100%) | 86.0 ± 22.3, *0-100* | 79.7 ± 26.3, *0-100* | 83.5 ± 22.2, *0-100* |
| Frequency (days/month) | 2.40 ± 3.42, *0-15.04* | 2.83 ± 3.84, *0-20* | 2.31 ± 3.33, *0-15.29* |
| Expected satiety (0-100%) | 42.3 ± 26.0, *0-100* | 59.1 ± 19.4, *0-100* | 47.1 ± 23.0, *0-100* |
| Healthiness (0-100%) | 36.7 ± 24.3, *0-100* | 46.5 ± 20.0, *0-100* | 35.9 ± 22.3, *0-94.1* |
| Estimated energy content (kcal) | 115 ± 50, *17-240* | 122 ± 45, 21*-240* | 110 ± 46, 0*-240* |
| Estimated energy density (0-100%) | 49.0 ± 24.5, *0-100* | 66.8 ± 17.2, *0-100* | 55.7 ± 22.4, *0-100* |
| Estimated price (USD) | 1.34 ± 0.96, *0.07-5* | 1.57 ± 1.11, *0-5* | 1.29 ± 0.99, *0.06-5* |
| Willingness to pay (USD) | 0.94 ± 0.96, *0-4.92* | 0.98 ± 1.09, *0-5* | 1.04 ± 0.95, *0-4.55* |

**Table S2.** Descriptive statistics for subjective ratings and WTP for foods across the three macronutrient categories from participants with overweight/obesity (OW/OB; n = 30).

| **Subjective Variable**  (Units or scale range) | **Carbohydrate**  M ± SD, *Range* | **Fat**  M ± SD, *Range* | **Combo**  M ± SD, *Range* |
| --- | --- | --- | --- |
| Liking (-100 to +100%) | 17.1 ± 32.4, *-90.4 to +100* | 17.6 ± 31.0, *-95.4 to +100* | 21.6 ± 30.4, *-86.9 to +100* |
| Familiarity (0-100%) | 80.3 ± 24.0, *0-100* | 75.7 ± 28.0, *0-100* | 78.6 ± 24.6, *0-100* |
| Frequency (days/month) | 2.28 ± 3.57, *0-20* | 3.00 ± 4.12, *0-20* | 2.05 ± 2.82, *0-10.81* |
| Expected satiety (0-100%) | 43.5 ± 24.3, *0-100* | 55.8 ± 18.4, *0-100* | 43.5 ± 21.0, *0-94.9* |
| Healthiness (0-100%) | 37.4 ± 23.5, *0-94.6* | 48.3 ± 18.0, *0-100* | 35.9 ± 20.2, *0-81.4* |
| Estimated energy content (kcal) | 128 ± 52, *18-240* | 121 ± 48, *0-240* | 122 ± 53, *0-240* |
| Estimated energy density (0-100%) | 50.3 ± 22.0, *0-100* | 62.1 ± 14.6, *18.1-100* | 53.2 ± 30.9, *0-100* |
| Estimated price (USD) | 1.51 ± 0.99, *0.06-5* | 1.82 ± 1.10, *0.04-5* | 1.50 ± 0.97, *0.06-5* |
| Willingness to pay (USD) | 1.07 ± 0.98, *0-4.38* | 1.15 ± 1.07, *0-5* | 1.09 ± 0.98, *0-4.89* |

**Table S3.** Descriptive statistics and unpaired, two-sample t-tests on the Dietary Fat and Free Sugar Short Questionnaire (DFS) subscores and internal state ratings across BMI groups.

| **DFS Score or Internal State**  (Scale range) | **HW**  M ± SD, *Range* | **OW/OB**  M ± SD, *Range* | **BMI Group Difference** |
| --- | --- | --- | --- |
| DFS fat (0-55) | 28.1 ± 5.7, *19-43* | 28.7 ± 6.0, *19-46* | t_(1,58)_ = 0.379, p = 0.706 |
| DFS sugar (0-45) | 18.7 ± 4.6, *11-29* | 18.2 ± 5.3, *9-32* | t_(1,58)_ = -0.346, p = 0.731 |
| DFS fat-sugar (0-30) | 13.6 ± 3.3, *7-20* | 12.7 ± 3.2, *9-22* | t_(1,58)_ = -1.143, p = 0.258 |
| Hunger (0-100%) | 71.5 ± 18.4, *0.1-97.4* | 67.1 ± 23.2, *0-99.4* | t_(1,58)_ = -0.819, p = 0.416 |
| Fullness (0-100%) | 23.9 ± 20.9, *0-84.7* | 23.2 ± 19.5, *0-66.0* | t_(1,58)_ = -0.122, p = 0.903 |
| Thirst (0-100%) | 69.4 ± 20.0, *25.4-100* | 73.2 ± 19.9, 20.0*-100* | t_(1,58)_ = 0.751, p = 0.456 |
| Desire to eat (0-100%) | 72.1 ± 17.3, *26.7-98.9* | 66.4 ± 18.4, *25.9-100* | t_(1,58)_ = -1.241, p = 0.220 |
| Potential to eat (0-100%) | 71.4 ± 16.5, *26.5-100* | 66.4 ± 14.3, 26.5*-98.0* | t_(1,58)_ = -1.257, p = 0.214 |

**Table S4.** BMI group interactions in the regressions between WTP, actual energy density (AED), and estimated energy density (EED), tested on averages per food across participants with HW versus OW/OB.

| **Variables** | **Foods Included** | **BMI Group Interaction** |
| --- | --- | --- |
| **Outcome: WTP** | All stimuli | $\beta$ = 0.104, p = 0.627 |
| **Predictors: AED** $\boldsymbol{\times}$ **BMI Group** | Carbohydrate items | $\beta$ = -0.073, p = 0.858 |
|  | Fat items | $\beta$ = 0.159, p = 0.690 |
|  | Combo items | $\beta$ = 0.351, p = 0.262 |
| **Outcome: WTP** | All stimuli | $\beta$ = 0.017, p = 0.511 |
| **Predictors: EED** $\boldsymbol{\times}$ **BMI Group** | Carbohydrate items | $\beta$ = 0.018, p = 0.736 |
|  | Fat items | $\beta$ = 0.011, p = 0.885 |
|  | Combo items | $\beta$ = 0.012, p = 0.838 |
| **Outcome: AED** | All stimuli | $\beta$ = -0.018, p = 0.480 |
| **Predictors: EED** $\boldsymbol{\times}$ **BMI Group** | Carbohydrate items | $\beta$ = -0.006, p = 0.897 |
|  | Fat items | $\beta$ = -0.028, p = 0.748 |
|  | Combo items | $\beta$ = -0.061, p = 0.206 |

**Table S5.** Correlations between actual energy density (AED) and characteristics and subjective ratings of foods in the combo category, tested on averages per food item across all participants.

| **Variable** | **Correlation with AED** |
| --- | --- |
| Liking | r^2^ = 0.392, p = 0.029 |
| Familiarity | r^2^ = 0.002, p = 0.884 |
| Frequency | r^2^ = 0.327, p = 0.052 |
| Healthiness | r^2^ = 0.637, p = 0.002 * |
| Expected satiety | r^2^ = 0.671, p = 0.001 * |
| Estimated energy content | r^2^ = 0.229, p = 0.115 |
| Estimated energy density | r^2^ = 0.587, p = 0.004 |
| Estimated price | r^2^ = 0.835, p < 0.001 * |
| Actual price | r^2^ = 0.859, p < 0.001 * |
| Volume | r^2^ = 0.703, p < 0.001 * |
| Visual area | r^2^ = 0.052, p = 0.478 |
| Fat content | r^2^ = 0.062, p = 0.436 |
| Carbohydrate content | r^2^ = 0.067, p = 0.417 |
| Protein content | r^2^ = 0.343, p = 0.046 |
| Sodium content | r^2^ = 0.550, p = 0.006 |

* p < 0.0033 after Bonferroni correction for the 15 correlations performed

**Table S6.** Testing if the relationships of food volume and expected satiety with actual energy density differ across pairwise comparisons of foods in the three macronutrient categories.

| **Variables** | **Pairwise Comparison** | **Macronutrient Category Difference** |
| --- | --- | --- |
| **Outcome: AED** | Carbohydrate versus Fat | t_(1,20)_ = 2.785, p = 0.011 * |
| **Predictors: Volume** $\boldsymbol{\times}$ **Macronutrient** | Carbohydrate versus Combo | t_(1,20)_ = 1.166, p = 0.257 |
|  | Fat versus Combo | t_(1,20)_ = 1.229, p = 0.233 |
| **Outcome: AED** | Carbohydrate versus Fat | t_(1,20)_ = 2.478, p = 0.022 |
| **Predictors: Expected Satiety** $\boldsymbol{\times}$ **Macronutrient** | Carbohydrate versus Combo | t_(1,20)_ = 2.45, p = 0.024 |
|  | Fat versus Combo | t_(1,20)_ = 2.211, p = 0.039 |

* p < 0.017 after Bonferroni correction for the 3 pairwise comparisons per combination of outcome and predictor variables.

**Table S7.** Participant characteristics of the independent cohort for testing the modified picture set with varying food portions (N = 22).

| **Characteristic** (Units) | **Mean ± SD, *Range*** |
| --- | --- |
| Sex | 6 Male, 16 Female |
| Age (yr) | 22.3 ± 6.6, *18-45* |
| Education (yr) | 14.5 ± 2.5, *12-20* |
| Race | 12 White, 1 Black/African American, 8 Asian, 1 More than one race |
| Ethnicity | 2 Hispanic or Latino, 20 Not Hispanic or Latino |
| Household income ^1^ | 5.2 ± 2.2, *1-8* |
| Height (m) | 1.68 ± 0.07, *1.58-1.79* |
| Weight (kg) | 62.35 ± 7.61, *48.10-77.25* |
| Body mass index (kg/m^2^) | 22.04 ± 1.72, *18.42-24.94* |
| Waist-hip ratio | 0.84 ± 0.07, *0.72-1.05* |
| Body fat (%) | 23.5 ± 7.1, *9.5-33.8* |

^1^ Household income was dummy coded from 1-8 according to 2018 US Census Bureau income percentiles.

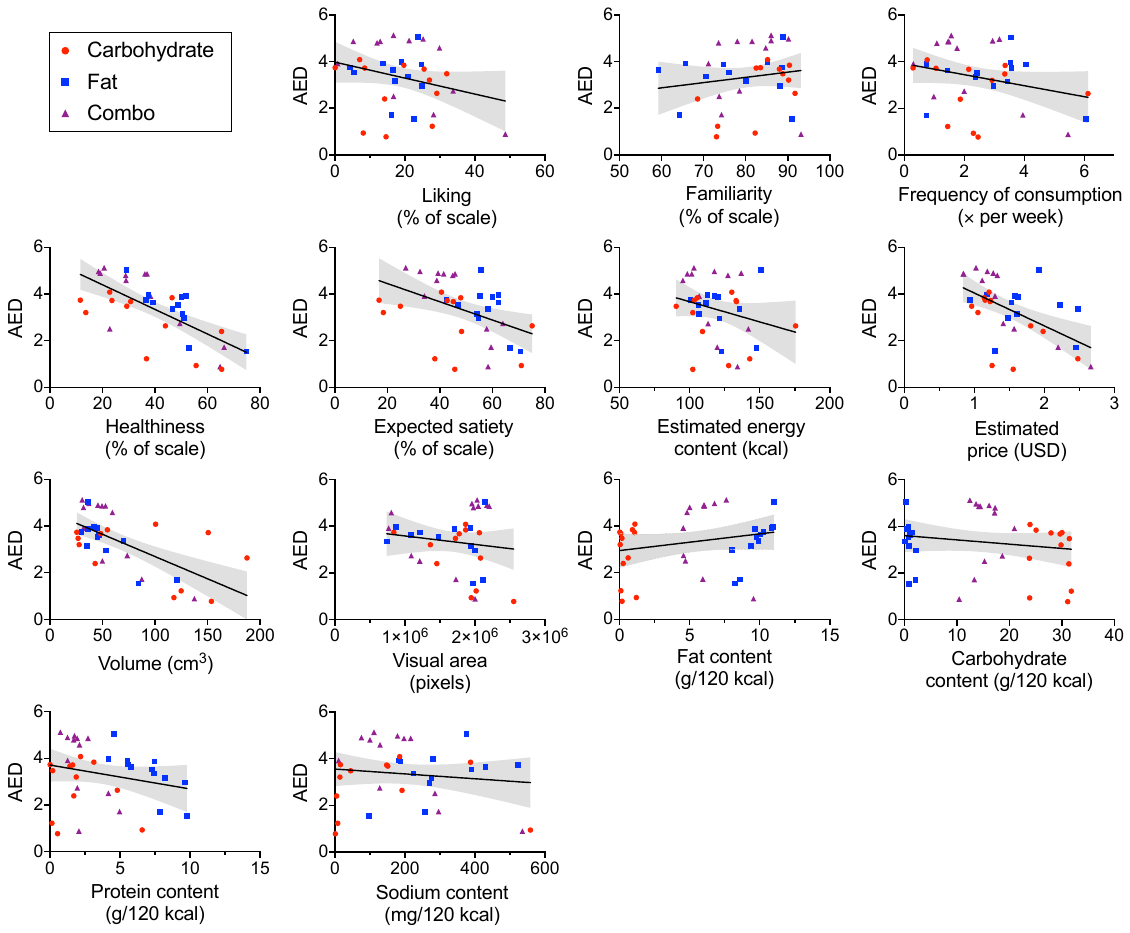

**Figure S1.** Fitted scatter plots comparing each food characteristic and subjective rating with actual energy density (AED, in g/120 kcal). Each data point represents a single food item from one of the three macronutrient categories (carbohydrate, fat, combo), with ratings averaged across all N = 60 participants (n = 30 HW and n = 30 OW/OB). Shading indicates 95% CI for the line of best fit.

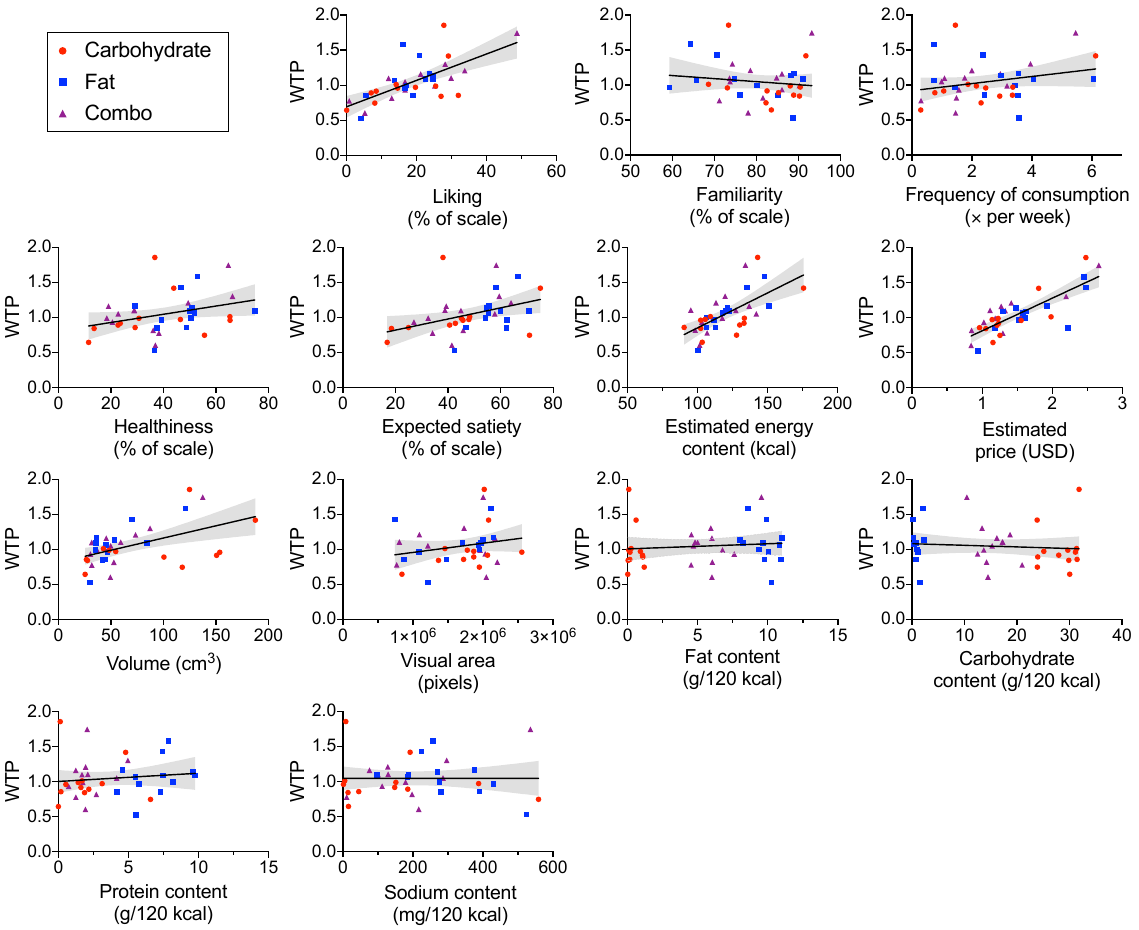

**Figure S2.** Fitted scatter plots comparing each food characteristic and subjective rating with willingness to pay (WTP, in USD). Each data point represents a single food item from one of the three macronutrient categories (carbohydrate, fat, combo), with ratings averaged across all N = 60 participants (n = 30 HW and n = 30 OW/OB). Shading indicates 95% CI for the line of best fit.

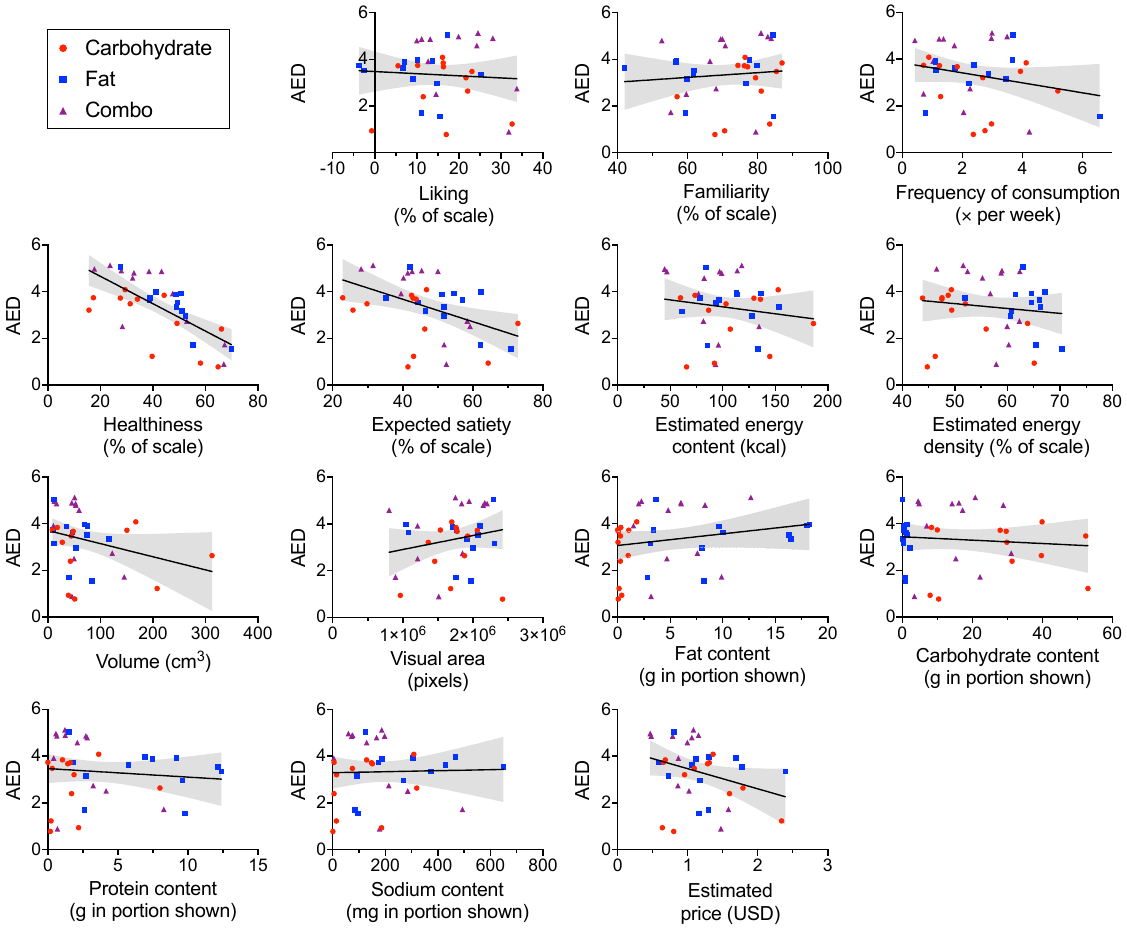

**Figure S3.** Fitted scatter plots comparing each characteristic and subjective rating with actual energy density (AED, in g/120 kcal) for foods pictured in varying portions (40, 120, 200 kcal). Each data point represents a single food item from one of the three macronutrient categories (carbohydrate, fat, combo), with ratings averaged across all N = 22 participants of the independent sample. Shading indicates 95% CI for the line of best fit.

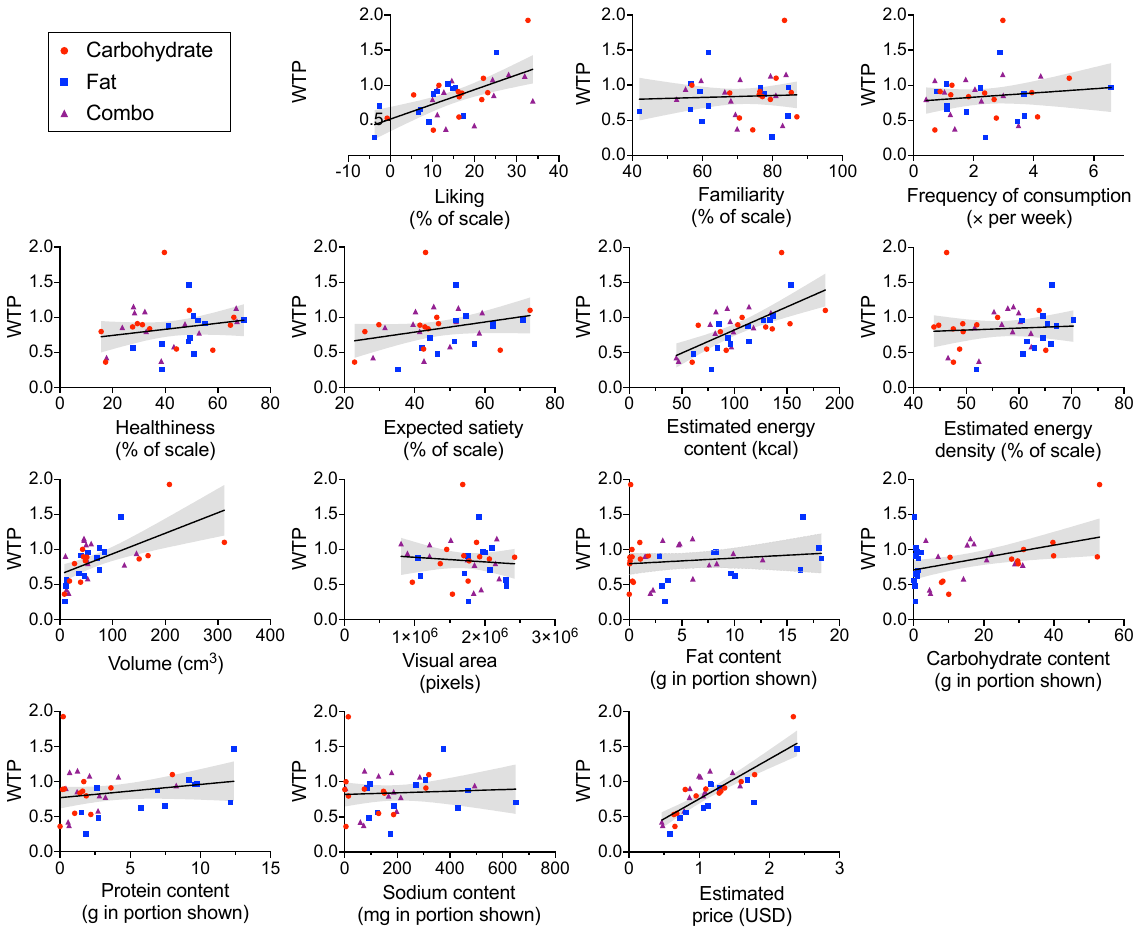

**Figure S4.** Fitted scatter plots comparing each characteristic and subjective rating with willingness to pay (WTP, in USD) for foods pictured in varying portions (40, 120, 200 kcal). Each data point represents a single food item from one of the three macronutrient categories (carbohydrate, fat, combo), with ratings averaged across all N = 22 participants of the independent sample. Shading indicates 95% CI for the line of best fit.
